# Muscle fatigue revisited - Insights from optically pumped magnetometers

**DOI:** 10.1101/2021.05.03.442396

**Authors:** Davide Sometti, Lorenzo Semeia, Hui Chen, Juergen Dax, Cornelius Kronlage, Sangyeob Baek, Milena Kirchgässner, Alyssa Romano, Johanna Heilos, Deborah Staber, Julia Oppold, Giulia Righetti, Thomas Middelmann, Christoph Braun, Philip Broser, Justus Marquetand

## Abstract

Muscle fatigue is well characterized electromyographically, nevertheless only information about summed potential differences is detectable. In contrast, recently developed quantum sensors optically pumped magnetometers (OPMs) offer the advantage of recording both the electrical current propagation in the muscle as well as its geometry, by measuring the magnetic field generated by the muscular action potentials. Magnetomyographic investigation of muscle fatigue is still lacking and it is an open question whether fatigue is characterized similarly in magnetomyography (MMG) compared to electromyography (EMG). Herein, we investigated the muscle fatigue during a 3×1-min strong isometric contraction of the rectus femoris muscle of 12 healthy subjects using simultaneous EMG-MMG (4-channel surface EMG and 4 OPM along the rectus femoris muscle).

Both EMG and MMG showed the characteristic frequency decrease in the signal magnitude during isometric contraction, which is typical for muscle fatigue. In addition, it was shown that the main part of this frequency decrease seems to occur in the circular component of the magnetic field around the muscle fibers and less longitudinally along the muscle fibers. Overall, these results show not only that magnetomyography is capable of reproducing the electromyographic standards in identifying muscular fatigue, but it also adds relevant information about the spatial characterization of the signal. Therefore, OPM-MMG offers new insights for the study of muscular activity and might serve as a new, supplementary neurophysiological method.

## Introduction

Muscle fatigue is the decrease in maximum force as a result of sustained or repetitive muscular activity [1]. The mechanisms that lead to muscle fatigue are manifold, but a major contribution is represented by fatigue of the innervating motoneuron and of the muscle cells themselves, as a result of accumulation of metabolites, such as lactate [2]. To measure or quantify muscle fatigue, surface electromyography (sEMG) is often used and well established. Here, muscle fatigue leads to a decrease in frequency of the electric muscular activity; the latter can be depicted in the form of power [3]. The electrical muscle activity corresponds to the muscle action potentials (MAP), of which the temporal and spatial sum can be measured using sEMG. MAP are action potentials at the neuromuscular endplate that propagate longitudinally along the muscle fibers as well as circularly via T-tubules around the muscle fibers. The resulting depolarization of the cell membrane triggers signal cascades, which cause an influx of calcium ions into the cell and an additional release of calcium ions from the sarcoplasmic reticulum, ultimately leading to contraction of the skeletal muscle [4].

### Electro- and Magnetomyography (EMG and MMG), optically pumped magnetometer (OPM)

As MAP propagate, Not only is an electrical potential difference generated, but also an electrical current along the direction of propagation, which, however, cannot be measured by sEMG. Furthermore, EMG is not sensitive to signal directions. These physical-technical limitations of electromyography (EMG) can be overcome by measuring the magnetic activity of the muscle with the so-called *magneto*myography (*M*MG, see also **Figure 1**): While EMG measures transmembrane potential differences (volts, *U or ΔV*), magnetomyography (MMG) maps indirectly the current flow (amperes, *I*) [5]. The basis for this is Biot-Savart’s law and Maxwell’s equations, which describe the following facts in a highly simplified way: Wherever an electric current flows, a magnetic field is generated, which can be measured using quantum sensors. Consequently, the MMG maps the current flow indirectly via the accompanying magnetic field. The idea that MMG could serve as a clinical neurophysiological diagnostic was proposed in the initial works on magnetometry in 1972 [6], but so far was not pursued due to the technical limitations of conventional magnetic sensors. Specifically, conventionally-used SQUID-sensors (superconducting quantum interference device) require cryogenic cooling to -269°C (4 Kelvin), and they lack ofspatial flexibility [7]. Recently, these limitations have been overcome by the development and improvement of optically pumped magnetometers (OPM) (for the physical background of OPM-MMG and OPM in biomagnetism, see e.g. [8], [9]). Due to the possibility of recording with the flexibly of arranged sensors and without cryogenic cooling, OPMs open up new opportunities to study muscles in complex anatomical situations [10]. Therefore, using MMG as an additional neurophysiological method to understand the electromagnetic processes of the muscle is now possible.

**Figure 1:**
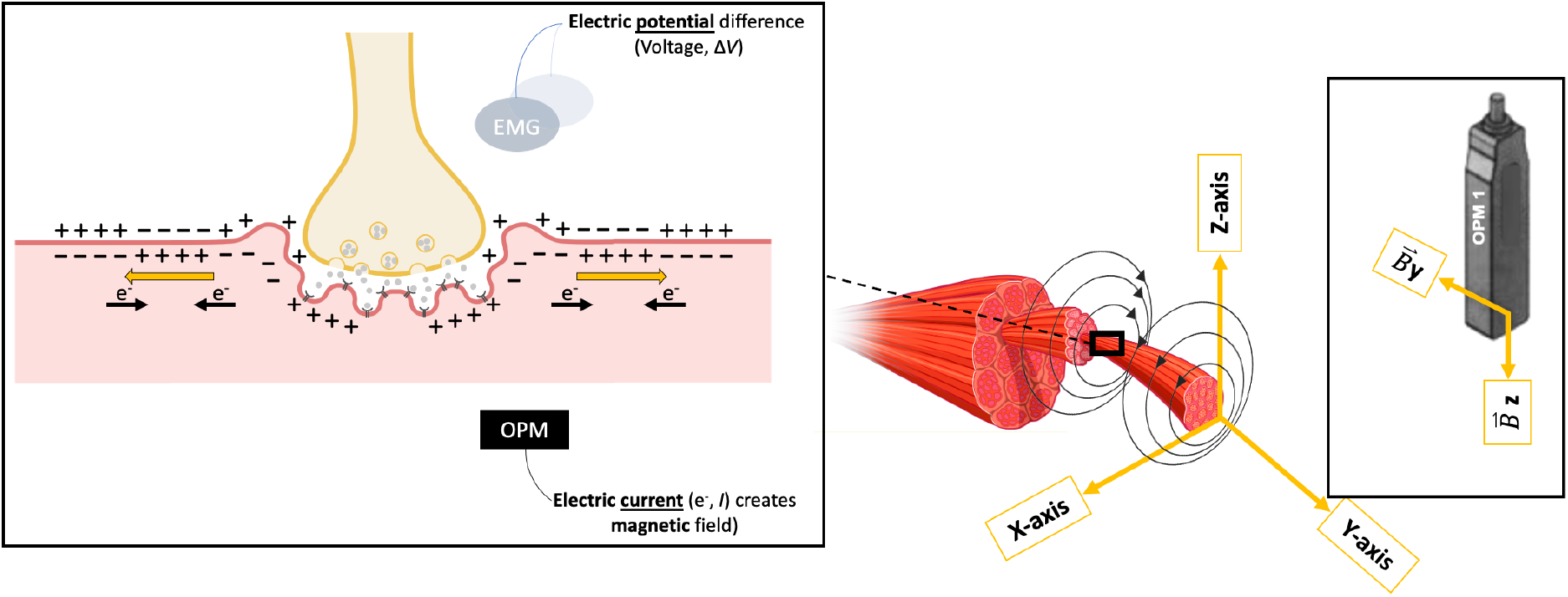
Basic principles of EMG and MMG. <u>Left</u>: Schematic drawing of a neuromuscular junction with two action potentials (yellow arrows) propagating from the neuromuscular junction along the muscle fiber. While the EMG measures potential differences (Δ*V*), the OPM-MMG indirectly measures the electric current (*I*) which coincides with a magnetic field. <u>Middle</u>: Simplified illustration of magnetic fields along muscle fibers. Magnetic fields have an orientation; thus, they have in principle vectorial components in all three geometrical directions (X, Y and Z). <u>Right</u>: OPM can measure the magnetic flux density 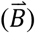 in two geometrical directions (Y and Z). Logically, by turning the OPM in a 90° angle, the 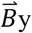 would be 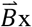.

### Approach towards muscle fatigue

After the first fundamental research and an understanding of the MAP by our group[8], [11], it seemed logical to investigate more complex situations of electrical muscle activity, i.e. muscle fatigue, and to compare it to the gold standard sEMG. In this context, we hypothesized that using OPM-MMG could not only offer equivalent results in terms of frequency decrease compared to sEMG, but could also yield new insights – such as the possibility of three-dimensional spatial signal acquisition as well as detecting vectorial field components is not possible using sEMG; the spatial properties of fatigue will be explored. New findings on muscle fatigue are highly relevant since muscle fatigue occurs in everyday life, in sports science[12], and in various neuromuscular diseases[13]. Therefore, we investigated the muscle fatigue strong isometric contraction of the rectus femoris muscle in 12 healthy subjects using simultaneous EMG-MMG.

## Methods

### Subjects

12 healthy subjects (6 males, 6 females) participated in the experiment (**Table 1**).

**Table 1:** Characteristics of the 12 participants (6 males, 6 females).

|  | **Average** | **Standard deviation** |
| --- | --- | --- |
| **Age** | 26.5 years | 2.9 years |
| **Weight** | 69.3 kg | 7.4 kg |
| **Height** | 176 cm | 7 cm |
| **Body mass index** | 22.3 kg/m<sup>2</sup> | 1.5 kg/m<sup>2</sup> |

**Table 2:** Pearson correlation coefficient (**r**), with a level of statistical significance expressed as p-value (**p**).

|  | <u>Center of gravity</u> |  |
| --- | --- | --- |
| | $r$ | $p$ |
| **OPM XX** | -0.69 | <0.0001 |
| **OPM XZ** | -0.37 | <0.0001 |
| **OPM YY** | -0.34 | <0.0001 |
| **OPM YZ** | -0.15 | 0.0002 |

**Table 3:** Pearson correlation coefficient (**r**), with a level of statistical significance expressed as p-value (**p**).

|  | <u>Center of gravity</u> |  |
| --- | --- | --- |
|  | *r* | *p* |
| **OPM X** | -0.66 | <0.0001 |
| **OPM Y** | -0.28 | <0.0001 |
| **EMG first** | -0.53 | <0.0001 |
| **EMG second** | -0.52 | <0.0001 |

The experiments were conducted at the MEG-Center of the University of Tübingen, Germany, in March 2021 and according to the standards by the World Medical Association. The subjects of this study were all authors of this publication, and gave their informed consent for their data to be published.

### Experimental setup

Prior to the experiment, for each subject the left rectus femoris muscle of each subject was imaged via high resolution muscle ultrasound (Mindray TE7, 14Mhz-linear probe) in order to determine the longitudinal axis of the muscle. After ultrasound imaging, the subject sat down on a comfortable chair inside a magnetically shielded room (Ak3b, VAC Vacuumschmelze, Hanau, Germany). Here, four paramagnetic surface electrodes (Conmed, Cleartrace^2^ MR-ECG-electrodes) and the 4 OPM (QZFM-gen-1.5, QuSpin Inc., Louisville, CO, USA) were placed in a distal to proximal order along a line and parallel to the longitudinal extend of the rectus femoris muscle (see also **Figure 2**). Additionally, one ground electrode was placed on the right shoulder and one reference channel was placed on the ipsilateral lateral knee. The OPM were placed 6-7cm proximal to the patella, at a 40 mm distance from each other and about 10-30 mm above the skin surface. The knee angle was controlled in all subjects at 150°, which was controlled visually using a goniometer.

**Figure 2:**
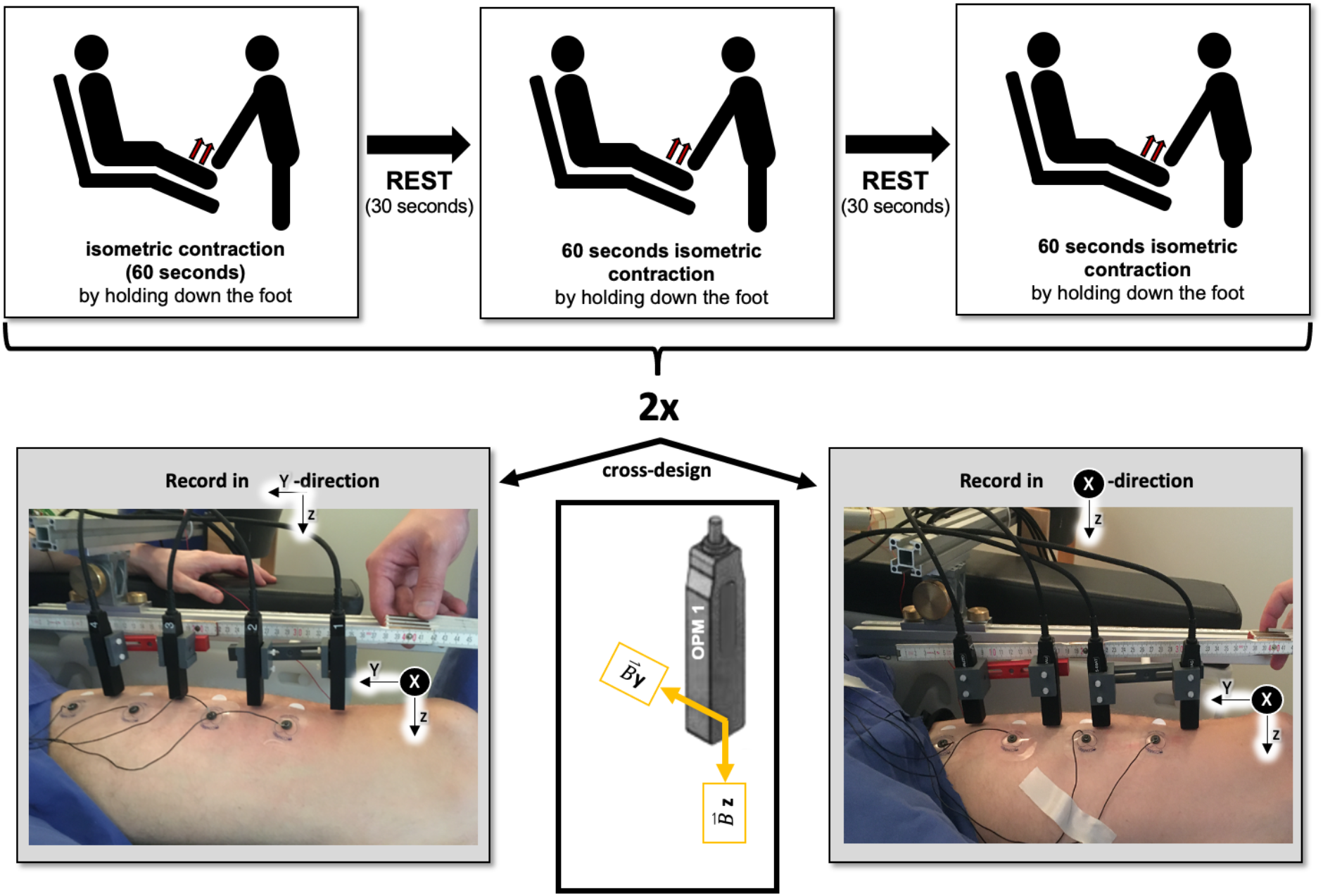
Illustration of the experimental EMG-MMG-setup. <u>Top</u>: Block design according to a 3×60 second isometric contraction with a 30-second rest in between. <u>Bottom</u>: After this, the OPM was rotated by 90° so that the X-direction could also be recorded, i.e., measurements were repeated according to 3×60-second isometric contraction with a break of 30 seconds each. According to a cross-design, 6 subjects were measured first in X-Z-and 6 subjects first in Y-Z-direction. In order to have a better overview of the relationship between spatial directions and measurable magnetic signals, an OPM including the measurable magnetic flux signals is depicted.

After this, the experiment was performed: The left ankle was pressed against the chair by a second person in the magnetically shielded room and the participants were asked to press against it with the strongest and most continuous force possible (**Figure 2**). This isometric contraction of the rectus femoris muscle took place three times in 60-second blocks, with a 30-second rest between each block. In order to be able to record the magnetic field vector components in all three spatial directions (X, Y and Z) by means of OPM-MMG and since the OPM used only record two vector components (in Y- and Z-direction), the OPM were rotated by 90° around their Z-axis to record also the vector component in the X-direction after the above-mentioned three 60-second blocks (**Figure 2, bottom**), and other 3 blocks were recorded with the updated set-up. This made it possible to acquire the magnetic flux signal in all three spatial directions. To avoid an unbalanced fatigue effects, the starting orientation of the OPM sensors was alternated across the subjects (cross-design). Following the same logic, we labeled the EMG recordings accordingly to the OPM orientation, thus respecting the same alternating pattern.

### Data acquisition

The analog output signals of the OPM System were recorded using the data acquisition electronics of an MEG System (CTF Omega 275, Coquitlam, BC, Canada) and the EEG channels of this MEG System were used to record the signals from the sEMG. The employed OPMs (QZFM-gen-1.5, QuSpin Inc., Louisville, CO, USA) were capable of measuring two components of the magnetic field vector: the y- and z-direction. They provided a magnetic field sensitivity in the order of 15 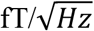 in a bandwidth of 3–135 Hz, an operating range below 200 nT and a dynamic range of a few nanotesla. To adapt to a non-zero magnetic background field, the sensors are equipped with internal compensation coils that can cancel magnetic background fields of up to 200 nT in the sensing hot rubidium vapor cell (vapor cell measuring 3 × 3 × 3 mm). The OPM system had an intrinsic delay of 3.8 ms which was corrected offline. The EMG signals were band-pass filtered with a Butterworth filter with an edge frequency set at 5 Hz (high pass) and 800 Hz (low pass). The data were post-hoc calibrated so that the magnetic fields and the electric potentials were in temporal synchrony.

### Data analysis

The analysis pipeline is illustrated in **Figure 3**. Data analysis was performed using MATLAB (MathWorks Inc., Natick, Massachusetts USA), and FieldTrip toolbox[14]. Continuously recorded EMG and OPM data were segmented in 3 trials of 60 seconds length, accordingly with the 3 isometric contraction sessions. In all the trials, one second of data-padding was left at the beginning and at the end of each trial in order to avoid edge artifact due to filtering processes. The signals were demeaned and filtered using a 10 Hz high-pass, zero-phase, 6^th^ order Butterworth infinite impulse response (IIR) filter. To suppress the powerline noise a band-stop (frequency ranges 48-52; 98-102; 148-152), zero-phase, 4^th^ order Butterworth IIR filter was applied. After the preprocessing, data was visually inspected, and 5 of the originally 12 subjects were discharged because of poor data quality due to technical problem during the recording, leaving 7 subjects with high quality data sets. Time-frequency analysis was performed using Morlet wavelet spectral power decomposition, with a Gaussian width of 15 (number of cycles) in order to maximize the frequency resolution. The time window of interest was set to 100 ms and the frequencies of interest ranged from 20 to 90 Hz in step of 2 Hz. We computed the magnitude of the signal taking the square root of the power, and finally we grand averaged the magnitude across the subjects. Since OPM can simultaneously record 2 orthogonal signal directions, Y and Z as well as X and Z (when the sensor is rotated of 90°), right before the grand average for each subject, we calculate the resultant component by applying the Pythagoras theorem so that:

**Figure 3:**
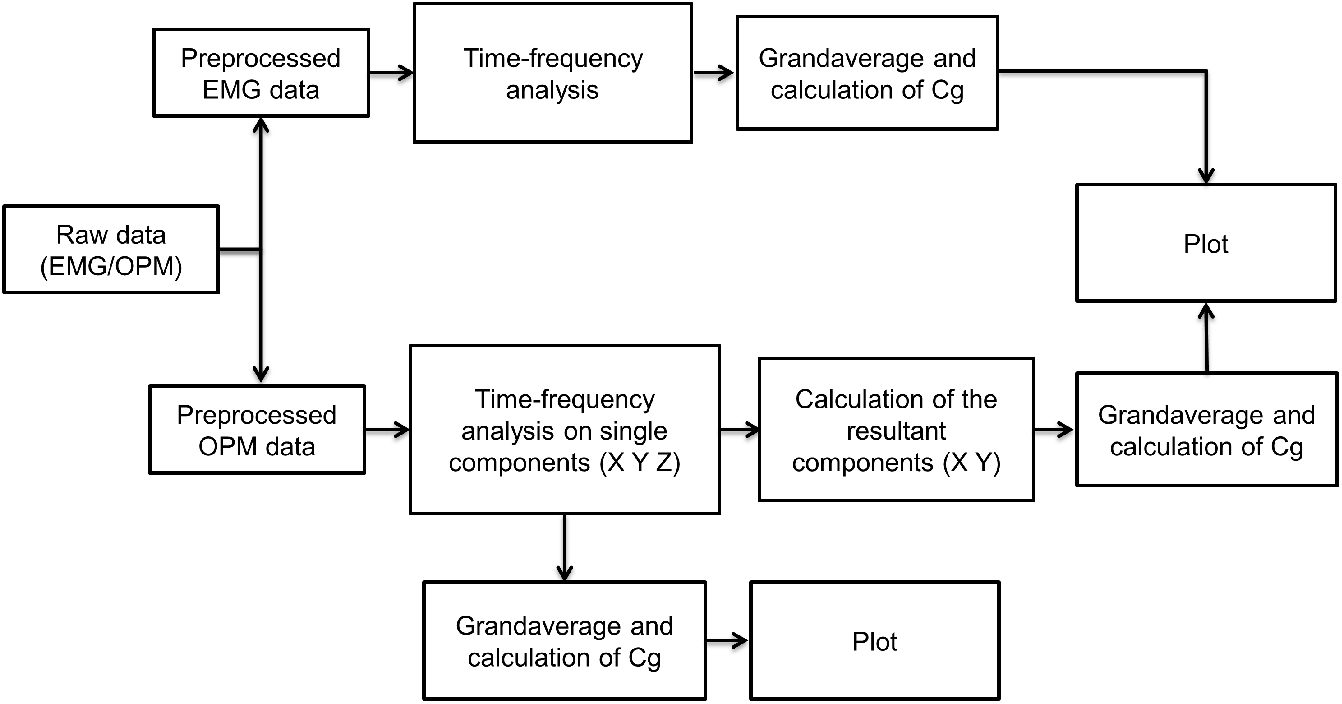
Illustration of the analysis pipeline: The time-frequency analysis of sEMG and separately OPM-MMG were calculated for each 60 second-block of isometric contraction and then averaged for OPM within a subject. These averages were then again averaged across all subjects to enable a group comparison between sEMG and OPM-MMG.

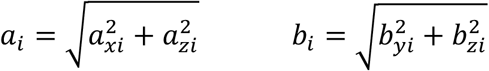

where *a* or *b* represent the magnitude of the frequency; *x, y*, and *z* are the 3 different directions for *i* OPM channels. Accordingly, *a*_*i*_ represents the sum of directions X and Z; *b*_*i*_ represents the sum of directions Y and Z. der In order to quantify the frequency decrease, the center of gravity (Cg) was calculated for each time points. Center of gravity represent the weighted average of the frequency over the entire frequency domain. Magnitude of the signal has been used here to weight the frequency so that:

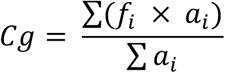

where *f* indicate the *i* frequency and *a* the respective magnitude. All the steps of the pipeline are illustrated in **Figure 3**.

## Results

### Time frequency analysis and spectral center of gravity in OPM-MMG

By rotating the OPM sensors by 90° (**Figure 2**), a total of 3 spatial directions (X, Y and Z) were recorded. In all the spatial directions, the OPM signal displayed the characteristic gradual frequency decrease in magnitude (**Figure 4**). Quantitatively, this decrease is noticeable by looking at the decrease of the center of mass of the frequency magnitude over time (**Figure 5**).

**Figure 4:**
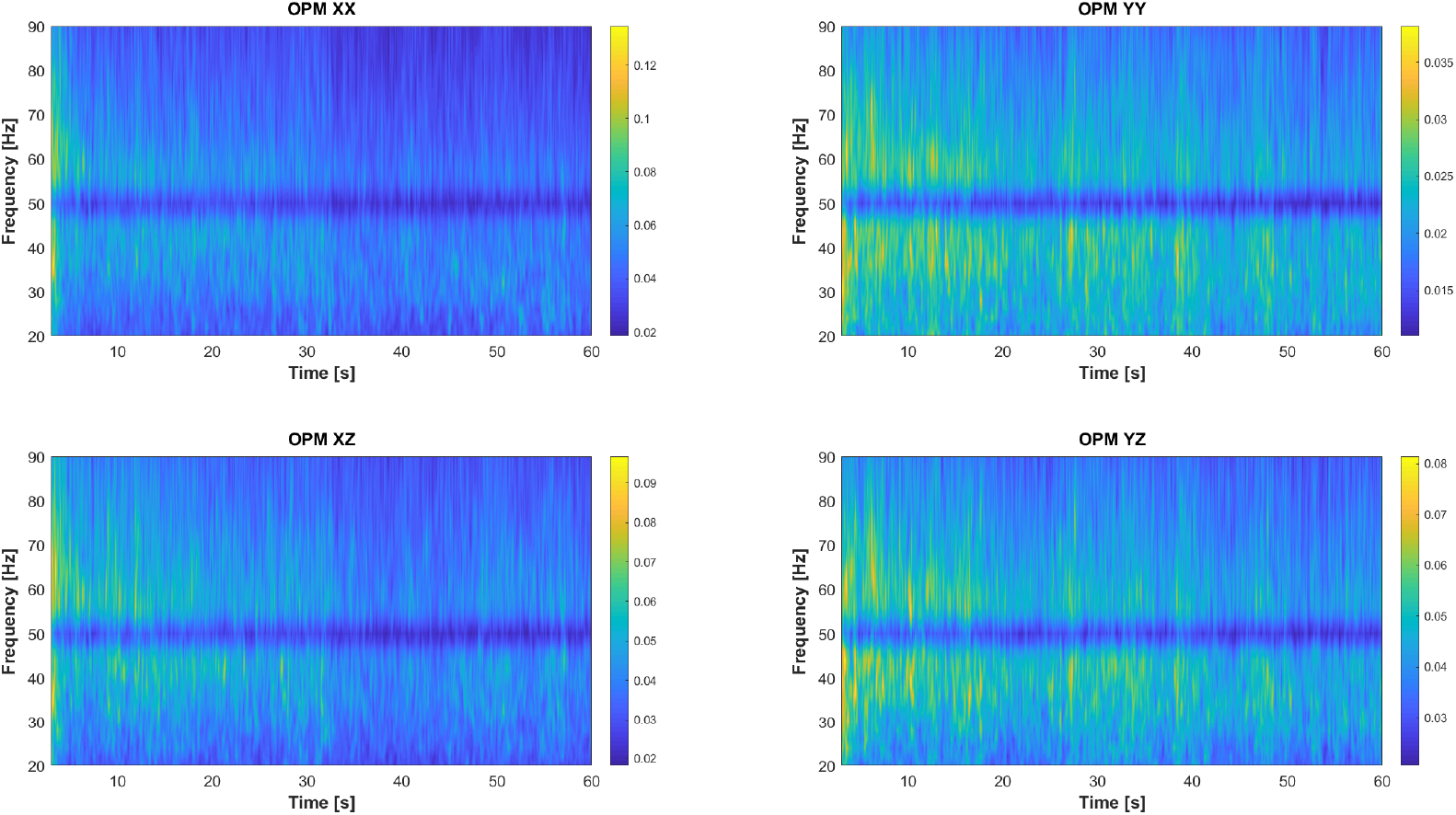
Time-frequency analysis of all OPM spatial directions X-Z (left) and Y-Z (right). Time on the x-axis from 0 to 60 seconds, frequency range on the y-axis from 20 to 90 Hertz, frequency magnitude in the colorscale with blue corresponding to low magnitude and yellow to high magnitude. To note, the band-stop filter was applied at 50 Hz for the powerline noise removal. The upper figure represents the magnitude of the frequency spectrum in time for the transversal (X) and longitudinal (Y) direction. Since the orthogonal spatial direction (Z) is always recorded3 by the OPM sensor, we analyzed the time frequency spectrum for the Z component during the transversal spatial direction recording (lower left) and from the longitudinal recording (lower right). For visualization purposes, the colorscale has not been matched between different figures, however, it is important to note how the intensity of the frequency spectrum for the transversal X direction (upper left) is one order of magnitude higher compared to the longitudinal Y direction (upper right). Visually, a decrease is appreciable in all the figure a frequency decrease in magnitude.

**Figure 5:**
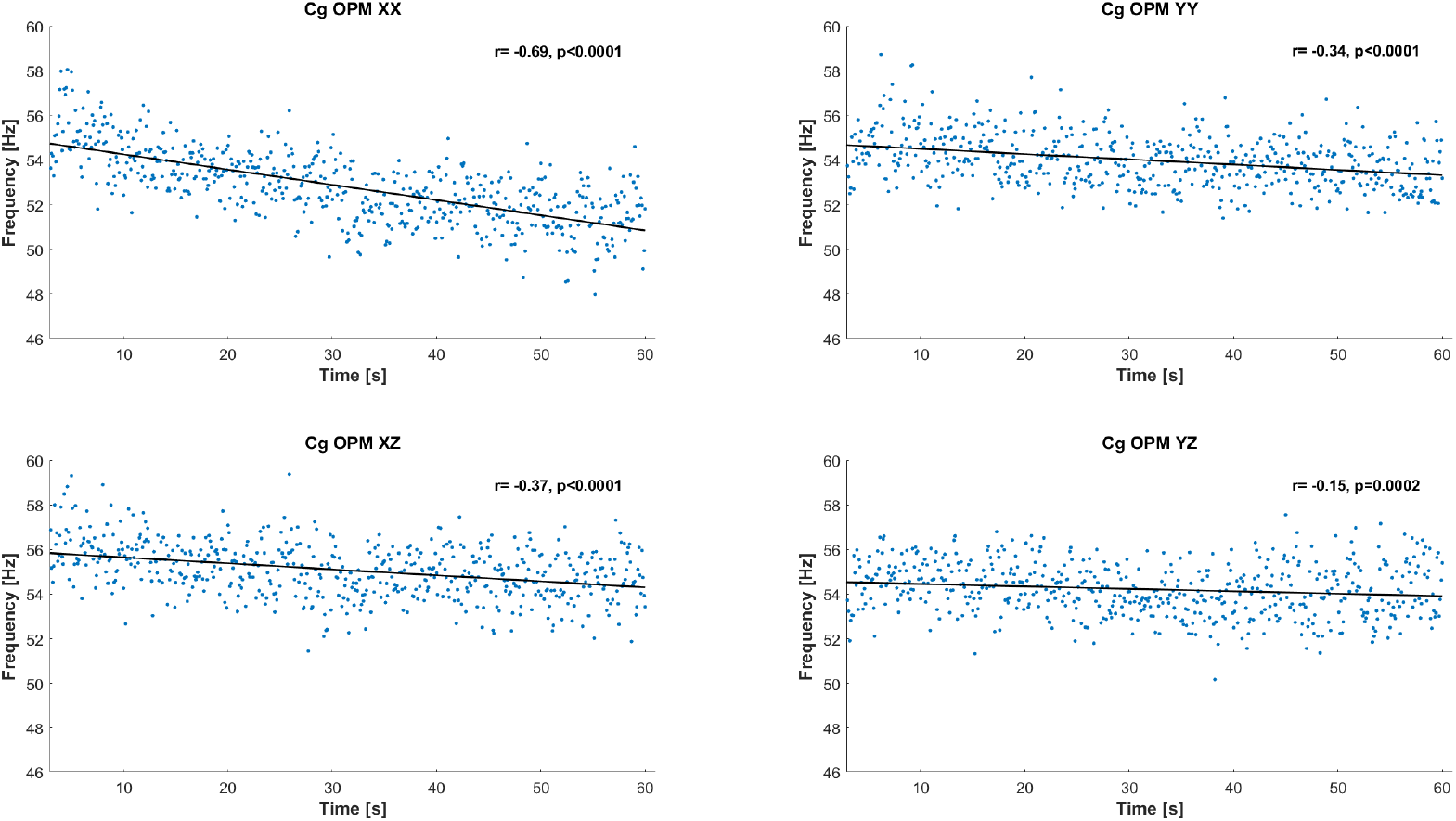
Spectral center of gravity (Cg) over time of all OPM spatial directions X-Z (left), Y-Z (right). Time in seconds on the x-axis, frequency in Hz on the y-axis. The spectral center of gravity provides a measure of the average height of the frequency in respect to its magnitude (blue dots maximum and minimum frequency at each time point, gray line represents a regression line). Here, the calculation was repeated for the entire trial at each time point. In all the OPM spatial recorded directions, the progressive decrease of the spectral center of gravity indicates that fatigue is occurring over time. The entity of this decrease is quantified by the Pearson correlation coefficient (**r**), with a level of statistical significance expressed as p-value (**p**).

### Comparison EMG-MMG

After applying the Pythagorean theorem on the orthogonal OPM direction X-Z and Y-Z, we were able to derive two resultant spatial components X and Y that we further compare with the EMG recording. Both sEMG and OPM-MMG displayed the characteristic spectrum decrease in magnitude over time (**Figure 6**). However, while the sEMG performance was constant between measurements, OPM-MMG performance differed depending on the orientation of the sensors. Despite both the orientation showed a frequency decrease over time, the transversal direction X, appeared to display higher sensitivity compared to the longitudinal direction Y (**Figure 7**).

**Figure 6:**
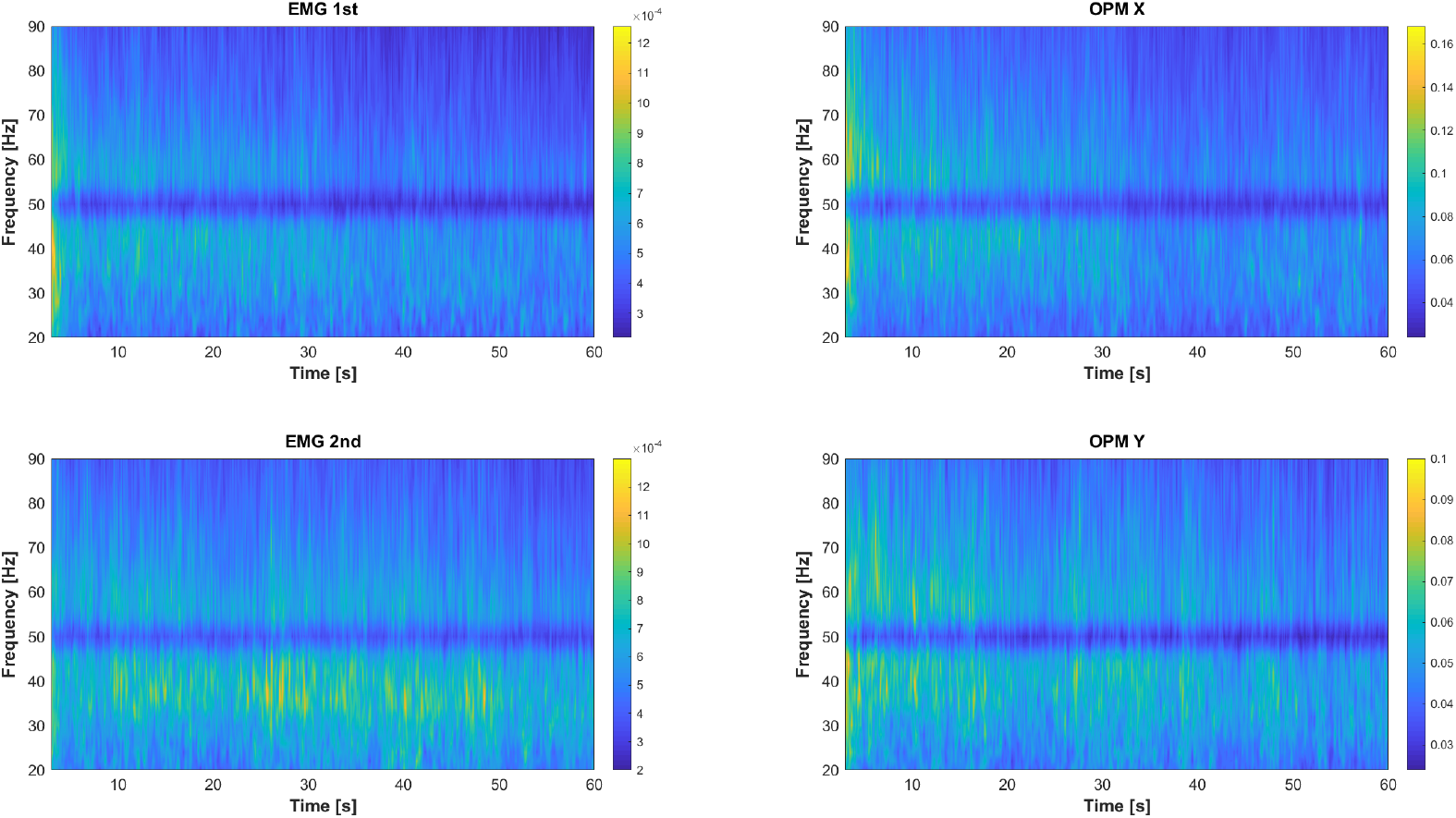
Time-frequency analysis EMG and OPM. Time on the x-axis from 0 to 60 seconds, frequency range on the y-axis from 20 to 90 Hz, frequency magnitude in the colorscale with blue corresponding to low magnitude and yellow to high magnitude. Different magnitude scaling depending on the different unit of measure (µV and nT). To note that the band-stop filter applied at 50 Hz for the powerline noise removal. Orthogonal OPM direction (X-Z and Y-Z) were summed using Pythagorean addition, leading to a resultant component that depending from the main spatial direction has been labeled **X** (upper right, 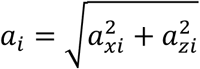) or **Y** (lower right, 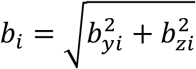). Both sEMG and OPM-MMG display the characteristic frequency decrease across time.

**Figure 7:**
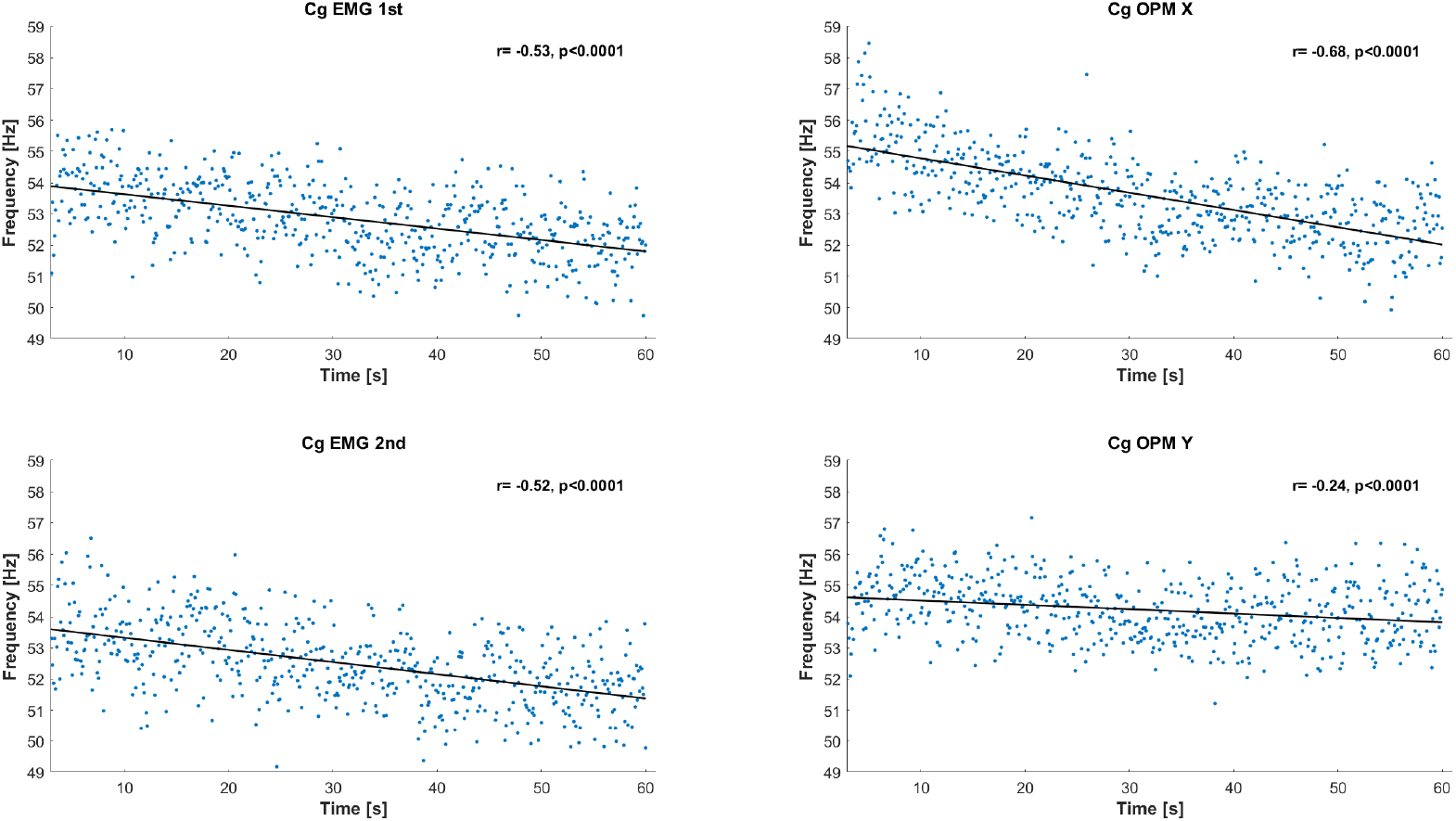
Spectral center of gravity (Cg) comparing EMG and OPM. Time in seconds on the x-axis, frequency in Hz on the y-axis (blue dots maximum and minimum frequency at each time point, gray line represents a regression line). A quantitative decrease of the spectral center of gravity over time is visible in both EMG (left) and MMG (right) signal. While this decrease is constant comparing different EMG recording, different OPMs orientation perform differently in depicting muscular fatigue, with higher sensitivity displayed by **X** (upper right, 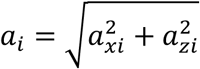) or **Y** (lower right, 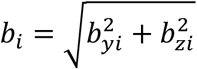).

## Discussion

This first investigation of the electro-magnetophysiology of muscle fatigue during isometric contraction of the rectus femoris muscle using simultaneous sEMG-MMG in the healthy participants reveals several new insights:

- OPM-MMG showed the characteristic frequency decrease of the power spectrum during muscle fatigue, which suggests that – as expected – the current flow frequency decreases during muscle contraction, just as the frequency of the electrical potential differences decreases over time.
- The main electromagnetic components of muscle fatigue seem to occur circular around the muscle fibers (X-direction) along the muscle fibers. A cautious interpretation of this novel result must be made (see below).
- The contactless OPM-MMG appears to produce at least equivalent results regarding time frequency analysis to the gold standard sEMG.
- We proof, that measuring muscle fatique with OPM-MMG in isometric contractions is feasible and that OPM can serve as a new neurophysiological method.

### Frequency decrease in power spectrum and geometry of magnetic flux signals in muscle fatigue

In both the gold standard sEMG and the OPM-MMG, we were able to detect the typical frequency decrease in the power spectrum of the duration of the isometric contraction, which is consistent with previous sEMG studies on muscle fatigue [15]–[17]. A challenging aspect of the analysis of magnetic signals is, the following circumstance: While the EMG measures electric potentials, where in the case of the muscle the special geometric orientation can be neglected, in the case of magnetic fields the geometric orientation has to be considered. This circumstance is both an advantage and disadvantage of the OPM-MMG: On the one hand, direction-selective magnetic signals of the muscle can be recorded, on the other hand, they cannot or can only be partially recorded if the OPM are not optimally aligned. In our study, this circumstance also became evident. The signals of the Z-direction were recorded twice, once in X-Z and Y-Z, which is shown in **Figure 4**. This finding appears conclusive when the underlying anatomy of the rectus femoris muscle is considered. The muscle fibers of the rectus femoris muscle do not run longitudinally in a parallel arrangement as in other muscles (e.g., biceps brachii muscle), but at an angle, which is called the pennation angle [18]. The pennation angle describes the angle between the muscle fibers of a muscle and its longitudinal axis. When the OPM are rotated from Y-Z to X-Z, the geometric orientation in the Z-direction to the oblique muscle fibers is also slightly changed, so that the signals are not acquired with the same intensity as before. In addition, by rotating the sensors, it is also possible to change the angle of the sensors themselves, which also affects the geometry.

However, the triaxial (X-Y-Z) signal acquisition of the OPM not only has the advantage in terms of spatial signal acquisition, but also a larger amount of data. Whereas EMG only ever captures one set of data – for example, a recording over 60 seconds of isometric contraction – OPMs capture at least two and therefore have twice as much data set as EMG. Potentially, this could be an advantage because more samples mean more information. Simply put, we collect twice as much information about a muscle with an OPM-MMG than with an EMG during a standard recording (at least in our setup).

This additional information was evident when considering the geometry of the main frequency decrease during isometric contraction. Here, we found that the main electromagnetic components of muscle fatigue occur radially (X-direction r -0.69) and not longitudinally (Y-direction r -0.34) along the muscle fibers. The interpretation of this result must be made cautiously, as several factors could potentially influence these findings and could not be controlled for in our setup: (1) Knowledge of the exact spatiotemporal propagation of the muscle action potential at the cellular level remains incompletely elucidated, i.e., we do not know how fast, how many muscle action potentials propagate from the neuromuscular endplate to where. (2) The geometry of the signal-generating muscle fibers and the positions of the signal-detecting OPM will necessarily differ from subject to subject, so that alterations in all three spatial directions are possible if the geometry is not fully known and controlled. In particular, the pennation angle, which decreases from distal to proximal in the rectus femoris muscle, may affect the results of the different spatial directions when averaged over 4 OPM.

Based on these findings, it can be summarized that MMG using OPM provides more information and insight into muscle physiology through geometric recording of muscle activity and fatigue, but also involves correspondingly more variables (geometry), whose control must be optimized and adapted. However, the problem of geometry with respect to MMG can be solved. For this – analog to magnetoencephalography – the position, properties and distance to the respective signal source needs to be known. One possibility would be, for example, to first perform OPM-MMG ex vivo on single muscle fibers and to empirically record the exact spatial propagation of single muscle action potentials and then to further investigate it in the context of larger muscle groups without significant pennation angle (e.g., M. biceps brachii).

### Strengths and limitations

Strengths of our study are the clear and hypothesis-based design and the simultaneous recording of two different modalities (EMG and MMG), so that a good comparability is possible. The study population was young, did not suffer from any neuromuscular disease and the gender distribution was balanced, so that there is sufficient probability that the normal muscle physiology could be recorded. Limitations exist mainly with respect to the difficulty of the suboptimal geometry of the muscle studied. The pennation angle of the rectus femoris muscle is not optimal, the fibers run pinnately and not longitudinally from distal to proximal, as is the case for example with the biceps brachii muscle. Since it is impossible to keep the penneation angle exactly the same between subjects, this could distort the signals. Furthermore, it was not possible to keep the distance between OPM and muscle always the same, because the anatomy of 12 different legs is always slightly different, an example would be the amount of subcutaneous fat. In addition, we also did not control muscle strength during the experiment, which is otherwise often measured in studies on muscle fatigue. However, due to our research questions, whether and how the frequency decrease is characterized, the assessment of muscle strength was dispensable.

## Conclusions

OPM are capable of recording muscle fatigue contactless and add relevant information about its spatial characterization, offering new insight in the study of muscular activity. Although geometrical situation of both the muscle and the alignment of the sensors must be considered, OPM-MMG might serve as a new, supplementary neurophysiological method.

## Declarations

### Ethics approval and consent to participate

All subjects consented to participate in the study.

### Consent for publication

All subjects consented that the anonymized data is published.

### Availability of data and materials

The anonymized data and materials are stored locally and any raw data from the statistical analysis can be made available on reasonable request.

## Authors’ contributions

JM and DS conceptualized and conducted the study, analyzed the data, drafted and revised the manuscript. TM provided and set up the OPMs, analyzed the data, and revised the manuscript. JD assisted in conducting the study and built components to make a simultaneous measurement possible. CK, HC, SB, MK, AR, JO, JH, DS, GR, LS assisted in conducting the study and revised the manuscript. CB, PB analyzed the data and revised the manuscript.

All authors read and approved the revised manuscript

## Acknowledgements

We thank Markus Siegel from the MEG-center Tuebingen for enabling the study and Nathalie Alexander from the Children’s Hospital of Eastern Switzerland for the intellectual support when designing the fatigue experiment. We finally thank the International Max Planck Research School for the Mechanisms of Mental Function and Dysfunction (IMPRS-MMFD) for supporting Lorenzo Semeia.

## Abbreviations

OPM: optically pumped magnetometer
EMG: electromyography
sEMG: surface electromyography
MMG: magnetomyography
OPM-MMG: optically pumped magnetometer magnetomyography
MEG: Magnetoencephalography
SQUID: superconducting quantum interference device

